# Underground Heterosis for Melons Yield

**DOI:** 10.1101/2021.03.04.434025

**Authors:** Asaf Dafna, Ilan Halperin, Elad Oren, Tal Isaacson, Galil Tzuri, Ayala Meir, Arthur A Schaffer, Joseph Burger, Yaakov Tadmor, Edward S. Buckler, Amit Gur

## Abstract

Heterosis, the superiority of hybrids over their parents, is a major genetic force associated with plant fitness and crop yield enhancement. Understanding and predicting heterosis is crucial for evolutionary biology, as well as for plant and animal breeding. We investigated root-mediated yield heterosis in melons (*Cucumis melo*) by characterizing common variety grafted onto 190 hybrid rootstocks resulting from crossing 20 diverse inbreds in a diallel-mating scheme. Hybrid rootstocks improved yield by more than 40% compared to their parents and the best hybrid outperformed the reference commercial variety by 65% under both optimal and minimal irrigation treatments. To characterize the genetics of the underground heterosis we conducted whole-genome re-sequencing of the 20 founder lines, and showed that parental genetic distance was no predictor for the level of heterosis. Through inference of the 190 hybrids genotypes from their parental genomes, followed by genome-wide association analysis, we mapped multiple root-mediated yield QTLs. The yield enhancement of the four best-performing hybrid rootstocks was validated in multiple experiments with four different scion varieties. While root biology is receiving increased attention, most of the research is conducted using plants not amenable to grafting and, as a result, it is difficult to separate root and shoot effects. Here, we use the rich genetic and genomic resources of *Cucumis melo*, where grafting is a common practice, to dissect a unique phenomenon of root-mediated yield heterosis, by directly evaluating in the field the contribution of the roots to fruit yield. Our grafting approach is inverted to the common roots genetics research path that focuses mainly on variation in root system architecture rather than the ultimate root-mediated whole-plant performance, and is a step towards discovery of candidate genes involved in root function and yield enhancement.

**Highlight:** We show that yield heterosis is significant in melon and controlled independently above and underground. Using common-scion grafting approach, we find that heritable rootstock-mediated variation in a diallel population is associated with substantial fruit yield heterosis.

## Introduction

About 10,000 years has passed since humans have shifted from hunter-gatherer to agricultural societies (Bellwood *et al.*, 2007). While agricultural productivity has evolved at an exponential scale since then, human population growth and climate changes form today substantial challenges to global food security (Godfray *et al.*, 2010; Wheeler and von Braun, 2013; Gerland *et al.*, 2014). Genetic improvement of crop plant yield is therefore more important than ever for addressing these challenges in a sustainable manner.

The challenge in genetic analysis of yield reflects the biological complexity of this trait, as yield is an outcome of the cumulative effects of multiple factors over time and across plant organs. From a genetic point of view, this complexity implies the action of multiple genes that interact with each other and with the environment and explains the low heritability calculated for yield in genetic studies. Another complexity associated with the genetic architecture of yield is the prominent non-additive variance component for this trait. This deviation from additivity — also known as heterosis or hybrid vigor — is a major driver for yield improvement in crop plants (East, 1908; Shull, 1908). The impact of heterosis on agriculture is wide, and is estimated to globally cause 15-30% yield increases (Duvick, 2001). This impact is best demonstrated in corn breeding, in which a continuous linear yield improvement is ongoing for almost a century following the introduction of hybrid corn in the 1930s (Duvick, 2001; Troyer, 2006).

Empirical data in various species have shown that diverse genetic, molecular and physiological mechanisms are likely to explain heterosis, but we are still lacking a unifying theory that enables us to explain and predict heterosis of fitness-related traits, including biomass, growth rate and reproductive success (Lippman and Zamir, 2007; Chen, 2013; Birchler, 2015; Vasseur *et al.*, 2019). Several genetic hypotheses have been proposed to explain heterosis: *i*) Dominance: cumulative genome-wide dominance complementation that masks deleterious effects of non-shared recessive alleles. *ii*) Overdominance: also known as single-gene heterosis, a synergistic outperformance of heterozygous alleles at the same locus (Krieger *et al.*, 2010). *iii*) Pseudo-overdominance: a case of dominance that resembles overdominance because two recessive loci that complement each other are tightly-linked in repulsion (Li *et al.*, 2015), and *iv*) Epistasis: multi-locus inter-allelic interactions (Yu *et al.*, 1997; Li *et al.*, 2001).

Next-generation sequencing (NGS) technologies and the growing availability of whole-genome assemblies provide new tools to study heterosis. There is an ongoing effort to further explore and explain the underlying genetics and molecular basis of heterosis in model and crop plants (Huang *et al.*, 2016; Li *et al.*, 2016; Seymour *et al.*, 2016; Yang *et al.*, 2017a,b).

In parallel to these genetic studies on heterosis, there is a growing effort to improve plant productivity and adaptation through partially overlooked factor—plant roots. The influence of root characteristics on whole-plant performance is shown in model and crop plants and therefore root research is important for advancing plant biology and for the future of agriculture (Meister *et al.*, 2014; Rogers and Benfey, 2015). The challenge in root research is obvious: roots are underground and therefore less accessible for phenotypic characterization. A major part of the research is consequently directed to the development of phenotyping methods for root-system architecture (RSA) (Zhu *et al.*, 2011; Topp *et al.*, 2013; Rogers *et al.*, 2016). Genetic studies on roots are mostly focused on RSA variation, followed by testing the link between RSA and whole-plant performance. QTLs for RSA traits were mapped in tomato (Ron *et al.*, 2013), soybean (Manavalan *et al.*, 2015), maize (Zurek *et al.*, 2015), rice (Zhao *et al.*, 2018) and other crop plants. In rice, a causative gene, *DEEPER ROOTING 1 (DRO1)*, affecting root growth angle was cloned and shown to affect yield under drought stress (Uga *et al.*, 2013). Manifestation of heterosis in root development was also characterized in several studies on wheat (Wang *et al.*, 2006) and maize (Paschold *et al.*, 2010). However, while these studies and others are using advanced technologies to phenotype and genetically characterize RSA traits, the direct functional link to whole-plant performance remains challenging due to the inability to separate root effects from shoot effects.

Grafting is a common practice in fruit trees and several vegetable crops (mainly *Cucurbitaceae* and *Solanaceae*). The ability to separate and re-combine root and shoot of different genotypes within or even across plant species has an increasing impact on plant research and agriculture (Gregory *et al.*, 2013; Goldschmidt, 2014; Albacete *et al.*, 2015). Grafting is an efficient tool to deliver tolerance to soil-borne pathogens or to improve abiotic-stress tolerance (e.g. drought, salinity), through the use of tolerant rootstocks. It also plays an important role in physiological and developmental studies focused on signal movement across plant organs (Lifschitz *et al.*, 2006; Omid *et al.*, 2007; Shalit-Kaneh *et al.*, 2019). However, to date, the advantage of this experimental tool for genetic analyses of root function and direct effect on whole-plant performance is very limited, as reflected by the few published studies on QTLs and rootstock traits (Estañ *et al.*, 2009; Gur *et al.*, 2011; Tandonnet *et al.*, 2018; Asins *et al.*, 2020).

Melon (*Cucumis melo*) is an economically important species of the *Cucurbitaceae* family. It is among the most important fleshy fruits for fresh consumption worldwide with 28 million tons produced globally in 2019 (http://faostat3.fao.org/). *Cucumis melo* is extremely diverse for phenotypic traits and melons are cultivated in nearly all of the warmer regions of the world. Alongside the rich genetic resources available, the melon genome sequence was completed in 2012 (Garcia-Mas *et al.*, 2012) providing a solid anchor for advanced genomic research including recent whole-genome resequencing of more than 1,000 diverse melon accessions (Zhao *et al.*, 2019).

In the current research we use grafting—which is a common commercial practice in melon and other cucurbit crops—to separate between roots and shoot effects in order to specifically investigate, using a diverse diallel population, the mode of inheritance and impact of roots on yield variation and heterosis in melon.

## Materials and Methods

### Plant Material

#### Core melon panel

This research is centered on a core set of 25 diverse melon accessions (**Sup. Table 1**) that were selected based on genotypic and phenotypic characterization of a broader GWAS panel. The core set includes representatives of the two cultivated sub-species and the different horticultural groups in melon as well as the broad phenotypic spectrum available for key traits, as previously described (Gur *et al.*, 2017).

#### Creation of diverse, 25-way, diallel population

A multi-allelic population of 300 F1 hybrids was built through a half-diallel crossing scheme between the 25 diverse founders (**Figure 1**). Plants of the 25 parents were grown and intercrossed in the greenhouse at Newe-Ya’ar during the fall of 2017. We defined two subsets within the 25 founders set, where the smaller sets completely overlapped by the sets above them, and each corresponds with a half-diallel population specifically derived from its composition: *HDA10* – 10 parental lines and 45 F1 hybrids and *HDA20* – 20 parental lines and 190 F1 hybrids (**Sup. Figure 1, Sup. Table 1**).

**Figure 1:**
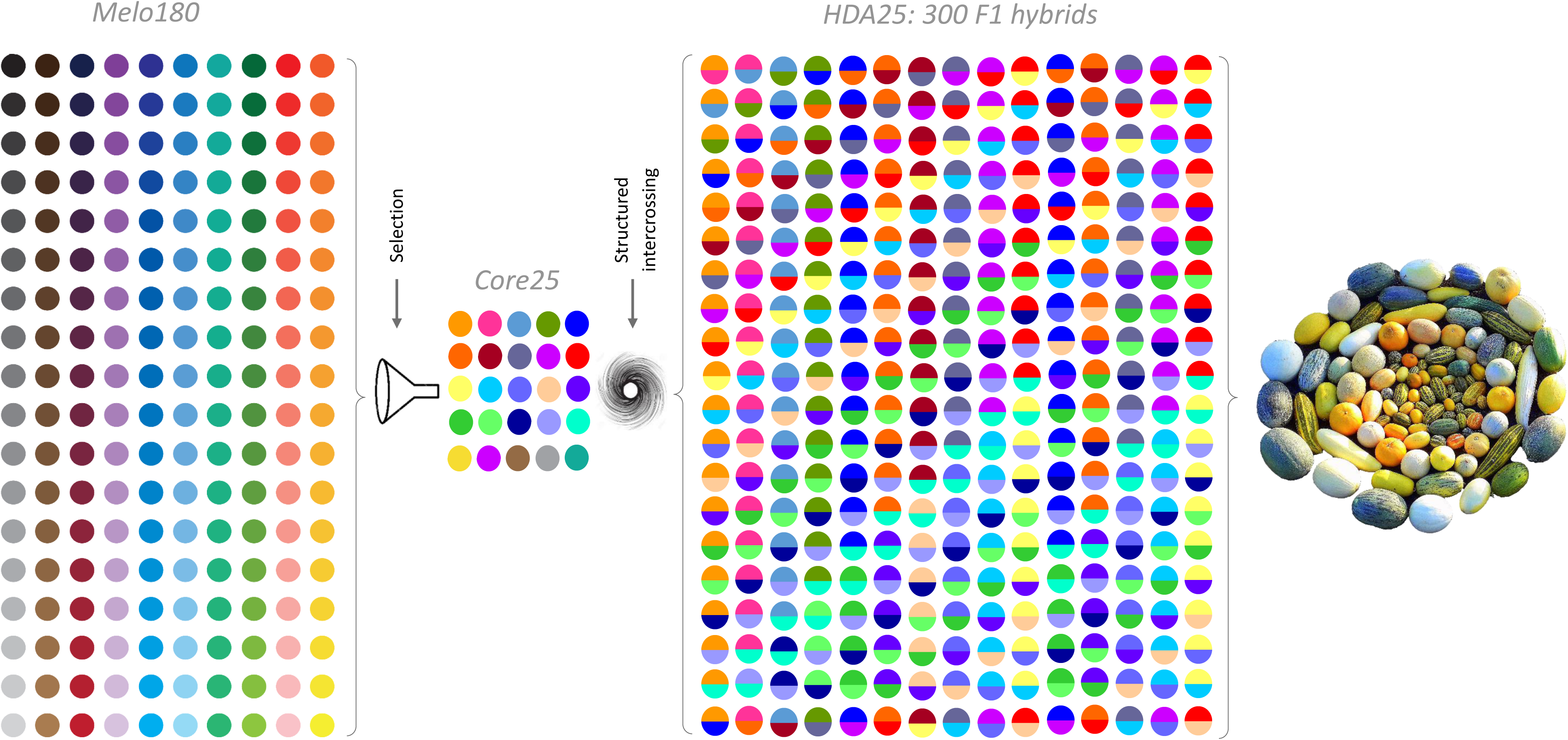
The path for development of the *HDA25* population. *Melo180* is a diverse collection (Gur *et al.*, 2017). *HDA25* is half-diallel population developed from the 25 core founders. On the right are representative mature fruits from the *HDA25* population.

### Field Trials

#### Non-grafted yield trials

Yield trials were performed during 2018, 2019 and 2020 spring-summer seasons, under standard growing conditions. Our main testing site is the open-field at the Newe Ya‘ar Research Center (32°43’05.4”N 35°10’47.7”E). Replicated trials consisted of three plots of five plants per plot in a randomized complete block design (RCBD). The standard planting density was 0.5 m between plants in a row and 1.90 m between beds. Selective harvest was performed at maturity of each genotype by going through the field 3 times a week over 4 weeks (mid-June to mid-July). All fruits from each ripe plot were harvested. Number of fruits (FN) and total fruit weight (Yield) per plot were collected at the field. Five representative ripe fruits were sampled from each plot for further analysis at the lab. Average fruit weight (AFW) was calculated on the sampled fruits as well as by dividing total yield by FN as measured at the field. Concentrations of total soluble solids (TSS, measured in degrees Bx) were measured on flesh samples from each of the five fruits separately, using hand-held refractometer (Atago A-10). Seeds were extracted from the sampled fruits, washed and dried and average seed weight was calculated from a sample of 50 seeds per replication (150 seeds per genotype).

#### Rootstock grafted yield trials

Each genotype (from either the *HDA10, HDA20*, parental lines or controls) was grafted as a rootstock with a common scion. Grafting for these large-scale experiments was performed in commercial nurseries (Hishtil - Ashkelon and Shorashim – Ein Habsor) under their standard grafting protocols. Shortly: rootstocks and scions were sown separately; approximately twenty-one days after sowing, seedlings from both rootstocks and scions were cut and grafted; plastic clips were used to attach the scion to the rootstock and promote efficient graft union development. Grafted plants were ready for transplanting in the field 7-10 days after grafting (**Sup. Figure 2**). The melon variety that was selected as the common scion for most parts of this project is ‘Glory’, a long shelf-life, high-yielding ‘Galia’-type variety. In addition to the good field-holding capacity of the mature fruits, this variety is also characterized by uniform fruit setting; both are critical attributes for this project, in order to allow performance of a single harvest of all yield. Each grafted entry was planted in five replications with five plants per replication (plot) in RCBD design. The standard planting density was 0.5 m between plants in a row and 1.90 m between beds. Drought stress treatment was applied by stopping the irrigation from start of fruit setting throughout the season until the harvest. Single non-selective harvest was performed when at least ~70% of the fruits were ripe and 95% reached their maximal size. In each plot, all fruits were harvested, counted and weighted for total yield calculation. Average fruit weight (AFW) was calculated by dividing the total fruit weight by the total number of fruits (FN) per plot. A sample of three representative ripe fruits was taken from each plot for total soluble solids (TSS) measurements performed on each fruit separately. Rootstock-mediated vegetative biomass was measured on grafted plants 56 days after transplanting (at the peak of female flowering and fruit setting) when most of the measured biomass is vegetative. The whole canopy of each plant was cut above ground level and fresh weight was measured.

### DNA preparation and genotyping

DNA was extracted using the GenElute™ Plant Genomic Miniprep Kit (Sigma-Aldrich, St. Louis, MO). DNA quality and quantification were determined using a Nanodrop spectrophotometer ND-1000 (Nanodrop Technologies, Wilmington, DE), electrophoresis on agarose gel (1.0%) and Qubit^®^ dsDNA BR Assay Kit (Life Technologies, Eugene, OR).

DNA of the 25 core accessions was shipped to the Genomic Diversity Facility at Cornell University (Ithaca, NY) for whole genome resequencing (WGS). Each sample was sequenced on an Illumina HiSeq 2000/2500 platform as 150 bp paired-end reads that were mapped to the C. *melo* reference genome DHL92 v4.0 (Ruggieri *et al.*, 2018), available at https://www.melonomics.net/melonomics.html#/download. SNP calling was carried out using the Broad Institute’s genome analysis toolkit (GATK ver. 3.7, McKenna et al. 2010), initially creating a separate genomic variant calling file (gVCF) for each individual detailing its polymorphism versus the reference genome, and later running a SNP discovery within the population. Initial SNP set was composed of ~9M SNPs that was filtered using TASSEL v.5.2.43 (Bradbury et al. 2007) for the following criteria: *i*) masking (as missing) scores with less than three reads per site, followed by the removal of sites with more than fifty percent missing data. *ii*) Minor allele frequency (MAF) >0.1. The final SNP set consisted of 4M SNPs. The whole-genome sequence alignment and derived HapMap from the 25 founders are now useful tools for detection of potential causative polymorphisms within candidate genes (Oren *et al.*, 2019)

### Statistical Analyses

JMP ver. 14.0.0 statistical package (SAS Institute, Cary, NC, USA) was used for statistical analyses. Mean comparisons were performed using the Fit Y by X function. GWA analysis was performed in TASSEL v.5.2.43 using the mixed-linear model (MLM) function. Distance matrix and Relatedness matrix of pairwise kinship (k matrix) were calculated in TASSEL from the filtered SNP dataset using the Centered_ IBS method (Endelman and Jannink, 2013). Stringent Bonferroni method was used to control for multiple comparisons in GWA. Best-parent Heterosis (BPH) was calculated as the deviation of the F1 hybrid from its better parent (F1-best-parent) and was expressed as absolute trait values or as Δ Percentage from best parent.

## Results

### Construction of diverse diallel population in melon

A primary resource for our genetic research on melon (*Cucumis melo*) is a diverse collection, composed of hundreds of accessions, which was built over the last 50 years in the Cucurbits Unit at Newe Ya‘ar (Burger *et al.*, 2009). We recently performed a Genome-Wide Association Study (GWAS) using 180 representative accessions and through comprehensive phenotyping and whole-genome GBS-based genotyping, demonstrated the effectiveness of this diversity panel for linkage-disequilibrium (LD) mapping of Mendelian fruit traits to candidate gene intervals (Gur *et al.*, 2017). Out of the 180 GWAS accessions, a core subset of 25 representative melon lines was selected based on integration of phenotypic and genotypic data; the core subset represents the two *Cucumis melo* subspecies and 11 horticultural groups. (**Sup. Table 1**, Gur *et al.*, 2017). Through structured intercrossing of the 25 lines in all possible combinations, we developed a half-diallel population (*HDA25*) composed of 300 F1 hybrids (**Figure 1**). This multi-allelic structure is a suitable design to characterize the mode-of-inheritance of traits, including general and specific combining abilities (GCA and SCA) patterns, and to perform GWAS on heterotic traits, such as yield.

### Above and underground yield heterosis in HDA10 population

To characterize yield variation and heterosis patterns, we first used a subset composed of 45 half-diallel F1 hybrids derived from intercrossing of 10 representative lines from our diverse collection (*HDA10*, **Sup. Table 1, Sup Figure 1**). These hybrids, placed alongside their parents, were tested in an open-field replicated yield trial during the summer of 2017. Half-diallel is a balanced design that reflects the same allelic composition and proportions in the F1 hybrids as in the set of parental lines and therefore allows informative general comparisons between the hybrids and inbreds sets, in addition to specific comparisons within hybrid groups (i.e. triads - hybrid and its two parents). In this experiment, hybrids fruit yield was on average 73% higher compared to their parental lines (**Figure 2a**). While variation in mode of inheritance of yield was observed across the 45 hybrid groups (**Figure 2b**), the superiority of hybrids over their parents was prevalent, with 13 F1 hybrids that showed significant best-parent heterosis (BPH). For example, *HDA10_005* is an F1 hybrid between a *C. callosus* line (P1, QME) and a *C. melo*, var *inodorous* line (P2, NA) that showed 90% best-parent yield heterosis (**Figure 2c**).

**Figure 2:**
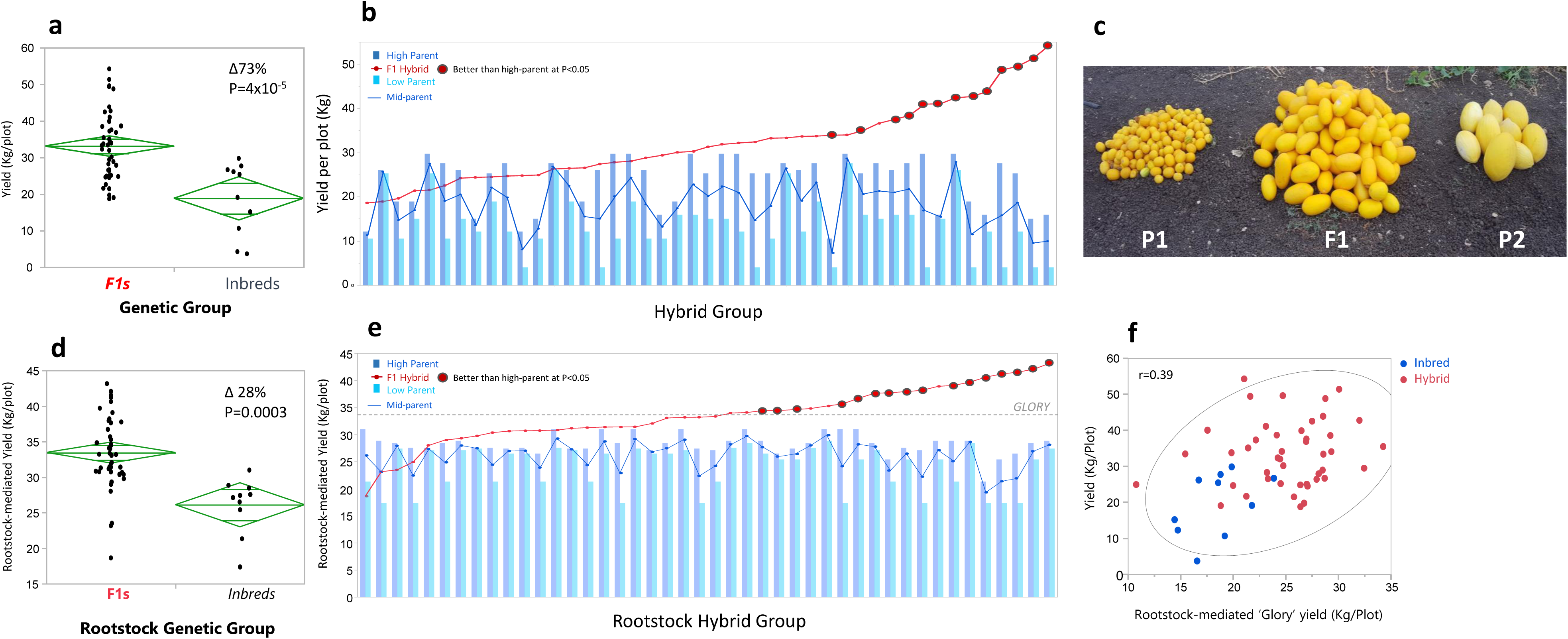
Yield heterosis across *HDA10* population (45 F1 hybrids and their 10 parental lines). **a)** Yield comparison between inbreds and F1s. **b)** Analysis of yield across 45 hybrid groups ordered in ascending manner by F1 yield. **c)** Example of heterotic hybrid (middle) alongside its parents. **d)** Root-mediated yield comparison between inbreds and F1s. **e)** Analysis of root-mediated yield across 45 hybrid groups ordered in ascending manner by F1 yield. **f)** Correlation between root-mediated yield (grafted) and yield of parallel genotypes in the non-grafted experiment, across the *HDA10* population.

In parallel to the conventional yield trial, we also tested whether yield variation and heterosis in melon can be derived from root effects *per se* and whether we can identify heritable variation for root-mediated effects. For this purpose, we took a grafting approach: the same germplasm set (45 *HDA10* hybrids + 10 Parents) were used as rootstocks grafted with a common commercial hybrid scion (‘Glory’, a long-shelf-life ‘Galia’-type hybrid). Such rootstock-grafting strategy allows us to eliminate the substantial aboveground variation across our germplasm and perform genetic analyses focused on the exclusive effect of the underground portion (roots) on yield. ‘Glory’ grafted on itself was used as control in this experiment. ‘Glory’ grafted with hybrid rootstocks yielded on average 28% more than parallel grafting with inbred rootstocks across the *HDA10* set (**Figure 2d**). Furthermore, most hybrid rootstocks across this set mediated higher yields as compared to their best-parents and 16 hybrid rootstocks showed significant BPH (**Figure 2e**). Overall, the proportion of yield variation explained by root-mediated genetic effects (broad-sense heritability) in this experiment was 40% (H^2^=0.40), a significant value that indicates a prominent contribution of roots to the yield variation. Moderate correlation was calculated between the rootstock-mediated yield and yield of the parallel *HDA10* hybrids and parental lines in the non-grafted experiment (r=0.39, **Figure 2f**), indicative of the independent aboveground variation components and the expected interactions between roots and scions.

### Rootstock-mediated yield heterosis across HDA20 population

Based on the positive results obtained on rootstock-mediated yield heterosis in the *HDA10* set, we extended the experiment and tested the wider *HDA20* set (190 half-diallel hybrids + 20 parents) as rootstocks grafted with the same common commercial hybrid, ‘Glory’, as scion. This set of 210 rootstock entries plus 2 controls (‘Glory’ grafted on itself and non-grafted ‘Glory’) was planted in replicated yield trial under optimal- and minimal-irrigation conditions (referred to as “Irrigated” and “Dry” herein, respectively). (**Sup. Figure 2a, b**). The Dry field yielded on average 30% less than the Irrigated and the correlation between the Dry and Irrigated trials was high (r=0.71, **Sup. Figure 2c, d**), and supported the significant genetic effect calculated for the root-mediated yield variation (H^2^=0.48). Further support for the significant genetic basis of the root effects is obtained from the correlations between the 2017 and 2018 grafted field experiments across the 55 *HDA10* genotypes (**Sup. Figure 2e, f**). Rootstock-mediated yield heterosis was apparent in both fields across *HDA20* population, with 38% (*P*=1.1×10^-8^) and 56% (*P*=1.8×10^-7^) average yield increase of hybrids compared to their inbred parents in the Irrigated and Dry fields, respectively (**Figure 3a, b**).

**Figure 3:**
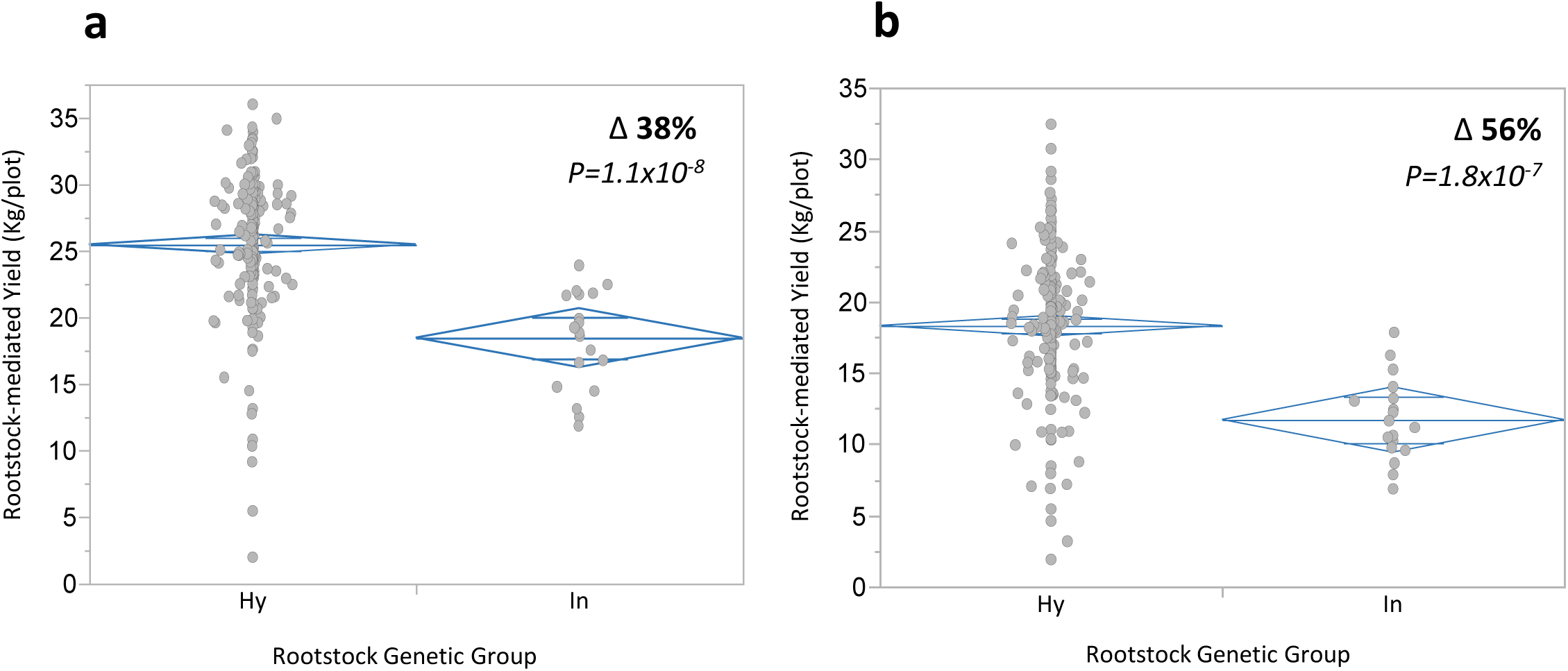
Root-mediated yield comparison between F1 hybrids and parental inbreds in the *HDA20* grafted rootstock yield trial. (a) Irrigated field. (b) Dry field.

The *HDA20* set can be viewed as 190 triads where each triad includes a hybrid and its two inbred parents; using this setup, we can define the mode of inheritance (additive and dominance components) within each triad, and draw patterns across the whole set. In this research, we use the stringent genetic definition of heterosis—the deviation of the hybrid from the high-parent (best-parent heterosis, BPH) —which is also the relevant definition from a breeding standpoint. The root-mediated yield of the 190 *HDA20* hybrids in the Irrigated and Dry experiments was, accordingly, partitioned to best-parent (BP) and heterotic (BPH) components (**Figure 4a, b**). A prevalent root-mediated yield BPH is evident, with 130 out of the 190 hybrids in the irrigated field showing a certain level of positive over-dominance, and 79 out of them displaying significant BPH (at *P*<0.05) and outperform their best-parent at an average of 55% (**Figure 4a**). The average BPH across all 190 hybrids was 26% (P=4.9×10^-30^) and 35% (P=1.2×10^-19^) in the Irrigated and Dry experiments, respectively. It is apparent from these results that (over)dominant deviation, a non-additive genetic component, is the major contributor to the root-mediated hybrid yield variation.

**Figure 4:**
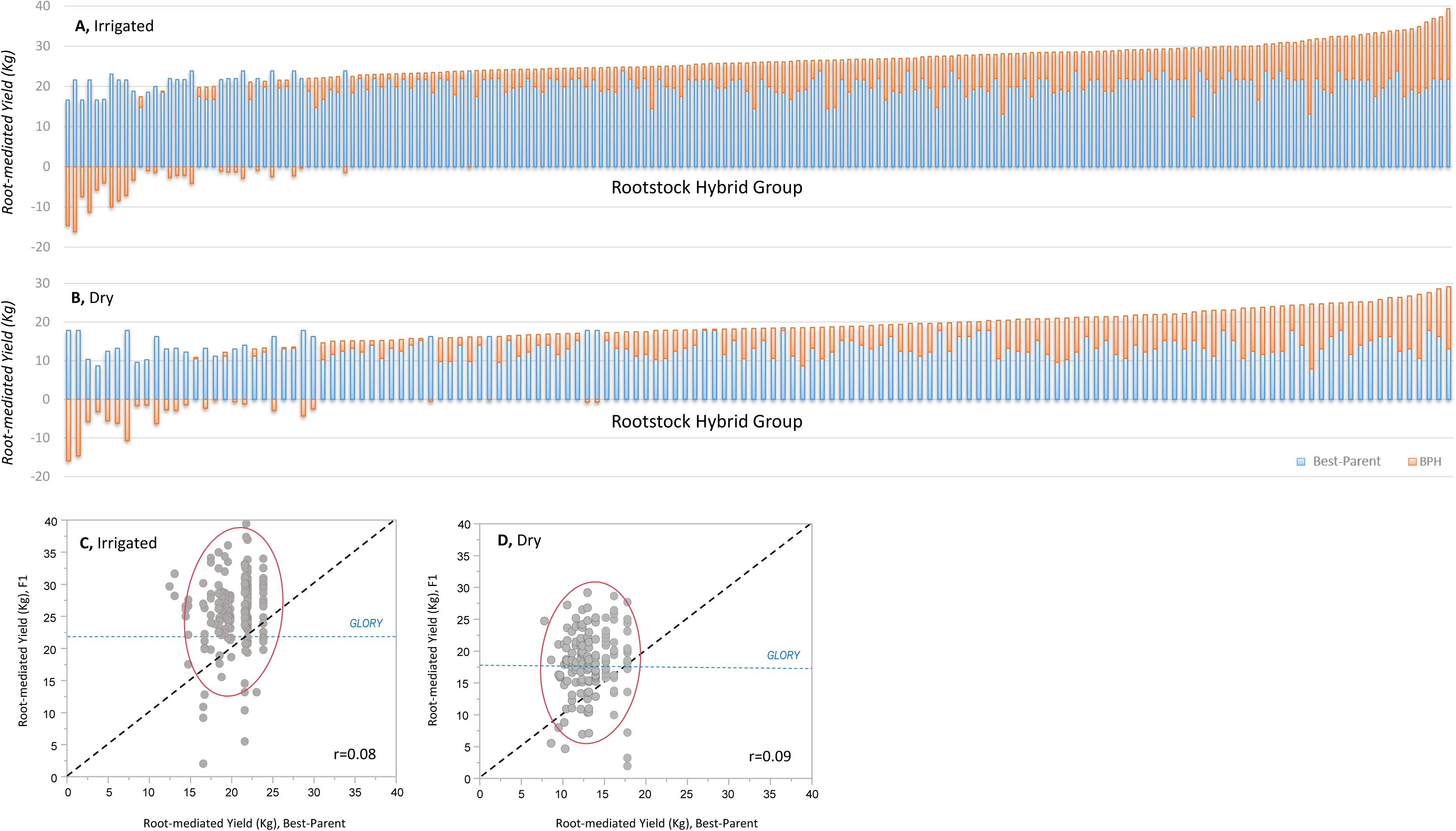
Partition of hybrids’ yield to parental and heterotic components. a and b) Yield of the 190 *HDA20* hybrids in the Irrigated and Dry fields, presented by its components: blue bars are the best-parent (BP) yield for each hybrid group, and orange bars represent the deviation of hybrid from best-parent (best-parent heterosis; BPH). Hybrids are ordered in an ascending manner by their yield. Negative orange bars reflect hybrids that are lower than their best-parent. c and d) correlations between root-mediated yield of best-parent and F1 hybrids across 190 *HDA20* triads. Dashed diagonal is x=y (BP=F1). Horizontal dashed blue lines are the yield of self-grafted ‘Glory’, the common scion variety.

Using the triads design, we could also test the broad relationship between parental and hybrid root-mediated yield performance across the diallel population. We show that there is no correlation between best-parents and hybrids root-mediated yield across the 190 hybrid triads in the Irrigated and Dry experiments (r=0.08 and r=0.09, respectively **Figure 4c, d**). This absence of correlation is supporting the observation that hybrid rootstock-mediated yield is independent of parental breeding value.

### Mode-of-inheritance of root-mediated yield compared to other melon traits

It was previously shown that heterosis is more prevalent in fitness-related, reproductive traits (Lu *et al.*, 2003; Rocha *et al.*, 2004; Semel *et al.*, 2006; Flint-Garcia *et al.*, 2009). We therefore collected data on additional traits in a non-grafted replicated experiment of this population (*HDA20*, 210 genotypes) and compared the general mode-of-inheritance between the root-mediated (grafted) yield and three seed- and fruit-related traits measured on non-grafted plants: average fruit weight (AFW), average seed weight (ASW) and flesh sweetness (total soluble solids, TSS). The comparison was performed by calculating the correlations between parental means and F1 hybrids across the 190 *HDA20* triads. While this correlation for root-mediated yield was essentially zero (r=0.01, **Figure 5a**), for AFW, ASW and TSS we found high positive correlations between hybrids and mid-parental performance (r=0.83, 0.92 and 0.80, respectively, **Figure 5b-d**). We also show that means of hybrids and mid-parents were not different in AFW, ASW and TSS of non-grafted plants, as compared with the 40% advantage of hybrids calculated for root-mediated yield (**Red triangles, Figure 5a-d**). Another visual way to demonstrate that non-additive, specific combining ability (SCA), is the prominent variation component of root-mediated yield across the *HDA20* population, is through the comparison of duplicated heat maps of the 20×20 half-diallel matrices of root-mediated yield (**Figure 5e**) and AFW (on the non-grafted experiment, **Figure 5f**). In these plots both dimensions are ordered by the average performance of each line across its hybrids (GCA) and the variation within rows or columns reflect the SCA. The uniform directional gradient apparent in AFW reflect the strong additive inheritance of this trait, while the mostly random distribution of high and low-performing hybrids in the root-mediated yield plot is indicative of non-additive inheritance. These analyses express the prominent additive component in the inheritance of the morphological and metabolic traits in melon, and demonstrate the fundamentally different mode of inheritance found for root-mediated yield.

**Figure 5:**
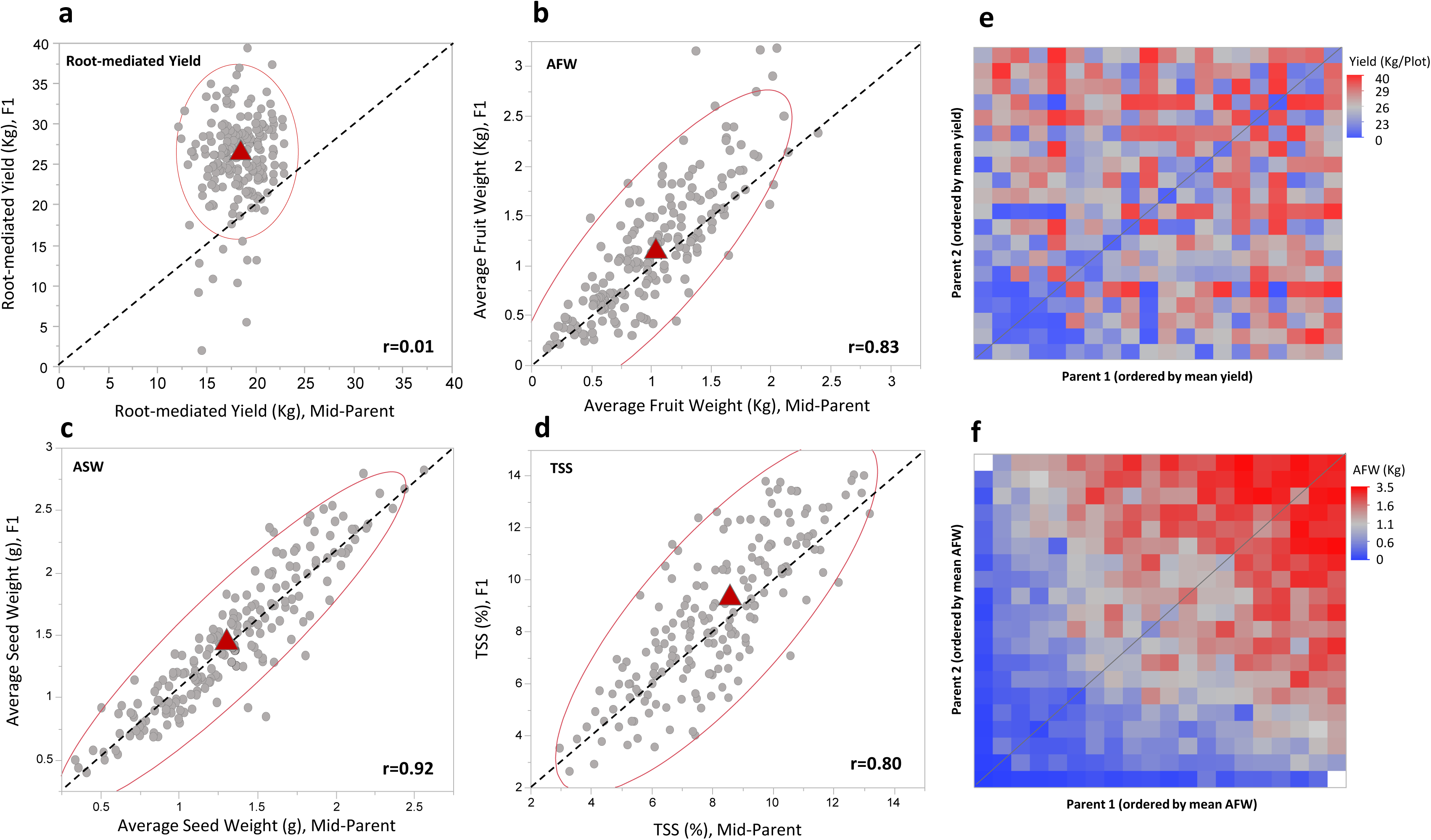
Correlations between mid-parent and F1 hybrid across 190 hybrid groups (*HDA20*). a) root-mediated yield (grafted). b) Average fruit weight (AFW, non-grafted) c) Average seed weight (ASW, non-grafted). d) Total soluble solids (TSS, non-grafted). Red triangles represent the averages of mid-parent and F1s. e) and f) present duplicated heat maps of the 20×20 half-diallel matrices for root-mediated yield (e) and for AFW (f). Both axes are ordered by parental GCA. Diagonals are the parents *per se* performance.

### Root-mediated effects on yield components and fruit quality traits in the HDA20 population

To describe further the nature of root-mediated effects across the *HDA20* population, we dissected the total fruit yield to its components—number of fruits per plot (FN) and average fruit weight (AFW). ‘Glory’ FN on a rootstock genotype-mean basis ranged between 11 and 30 fruits per plot and AFW range was 0.70-1.20 Kg/fruit. Surprisingly, both FN and AFW showed significant positive correlations with yield in the Dry and Irrigated experiments and accordingly were also positively correlated with each other (**Sup. Figure 3**). This pattern of yield variation and relation between its components is in contrast to the common negative tradeoff observed between FN and AFW across natural melon diversity, as we show in our non-grafted *HDA10* population (**Sup. Figure 4**). To assess the root-mediated effects on ‘Glory’ fruit quality, we also measured total soluble solids (TSS) on 2,100 fruits (10 fruits per genotype) across the grafted *HDA20* population in the Irrigated experiment. TSS is highly correlated with sugars content in the fruit flesh, which is a major determinant of melon fruit quality. The effect of rootstock genotype on TSS variation was not significant (H^2^=0.07) and accordingly was not correlated with the wide variation and high heritability of this trait across the *HDA20* population in non-grafted plants (**Sup. Figure 5**). Taken together, we find that high-yielding rootstocks are associated with more fruits, which are also larger on average, and these effects on yield and its components were not associated with any compensatory effect on fruit sugar content.

### Potential Predictors of root-mediated yield heterosis

The significance of heterosis, as shown above, in explaining hybrid root-mediated yield variation in melon, is providing an incentive to explore the genetic basis and underlying genes for this unique phenomenon and to develop predictive tools for effective breeding of heterotic yield-promoting rootstocks.

#### Root-mediated canopy biomass

We started by testing a potential simple phenotypic predictor. Using the same common-scion grafting setup, we measured root-mediated variation in plants canopy biomass across the *HDA20* set, and tested whether it is correlated with the root-mediated fruit yield variation. The rationale is that canopy vigor (biomass) is an easy-to-measure trait that can be phenotyped in high-throughput and cost-effective manner using remote-sensing technologies. While we also found heterosis for root-mediated plant vegetative biomass (**Figure 6a**), this trait is shown to be a poor predictor and explained only 3% of the root-mediated yield variation (**Figure 6b**).

**Figure 6:**
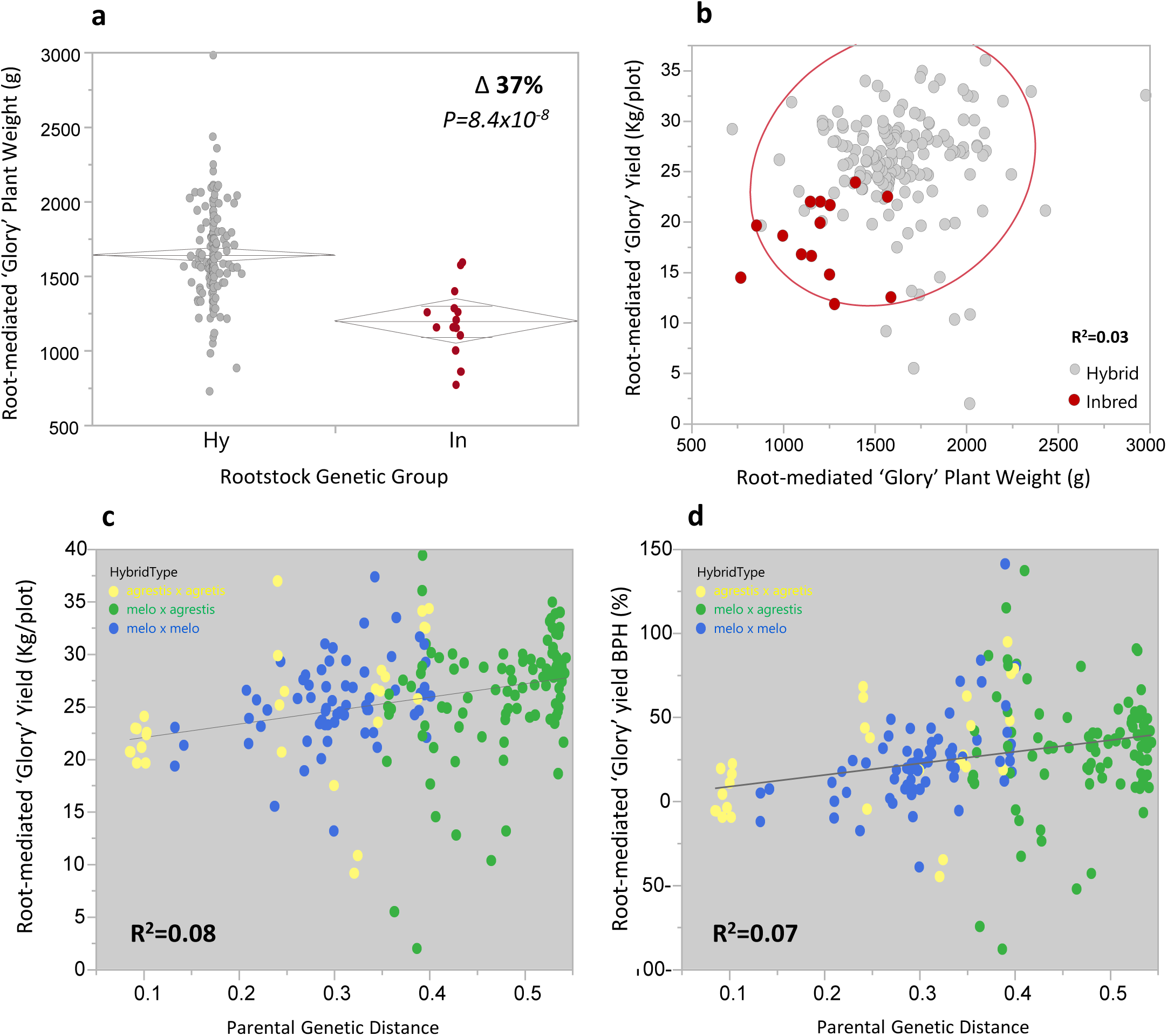
Potential predictors of hybrid root-mediated yield. a) Comparison of root-mediated young-plant vegetative biomass between *HDA20* hybrids and their inbred parents b) Correlation between root-mediated ‘Glory’ plant biomass and root-mediated ‘Glory fruit yield, across 156 hybrids + 13 inbred parents. c) Correlation between parental genetic distance and root-mediated yield, across 190 *HDA20* hybrids. d) Correlation between parental genetic distance and root-mediated yield BPH, across 190 *HDA20* hybrids.

#### Parental genetic distance

To test potential genetic predictors for root-mediated hybrid yield we conducted whole-genome re-sequencing of the 25 founder lines and extracted ~4,000,000 informative SNPs that describe the genetic variation. We show that parental genetic distance, which correspond with level of hetrozygosity at the F1, is also a poor predictor and explained only 8% of the root-mediated yield variation and 7% of BPH variation across the 190 *HDA20* F1 hybrids (**Figure 6c, d**). Accordingly, the type of the hybrid (*melo* and *agrestis*, inter or intra sub-specific) was also not predictive for rootstock performance. This lack of correlation between parental genetic distance or taxonomic classification and root-mediated hybrid yield may suggest that the yield heterosis is not confounded with relatedness or population structure and that there is a good chance of identifying specific loci significantly associated with this trait.

#### Root-mediated yield QTLs

To perform genome-wide association (GWA) analysis, we inferred the complete genotype (composed of 4,000,000 informative SNPs) for each of the 190 *HDA20* F1 hybrids, from their 20 parental genomes. We then used a filtered subset of 400K uniformly spaced SNPs (at parental minor allele frequency (MAF)>0.25) for the GWA analyses. The complex genetic nature of root-mediated yield variation is supported by multiple significant associations that were identified across the genome (**Figure 7**). On the irrigated experiment, we find significant SNPs on all chromosomes, and seven QTLs (on six chromosomes) are also common to the Dry experiment (**Figure 7b**). Allelic effects of two QTLs (q.RMY3.1 and q.RMY6.1) that were common to both environments are shown in **Figures 7c, d**. Both display heterotic inheritance, as the heterozygotes are associated with significant yield increase compared to homozygote genotypes in each SNP. While independently q.RMY3.1 and q.RMY6.1 explained 23%-25% (Dry, Irrigated) and 22%-28% (Dry, Irrigated) of the genetic variation, respectively (**Figure 7c, d**), joint haplotype of these SNPs significantly improved the model and explained 36%-37% of the variation. F1 hybrids that are heterozygotes in both QTLs are associated with higher root-mediated yield compared to those that are heterozygotes at one locus or other homozygote combinations (**Figure 7e**). The double heterozygote haplotype is associated with 16% and 14% root-mediated yield increase over the *HDA20* population mean, in the dry and irrigated fields, respectively. This effect reflects the estimated response to selection of favorable genotypes at these loci.

**Figure 7:**
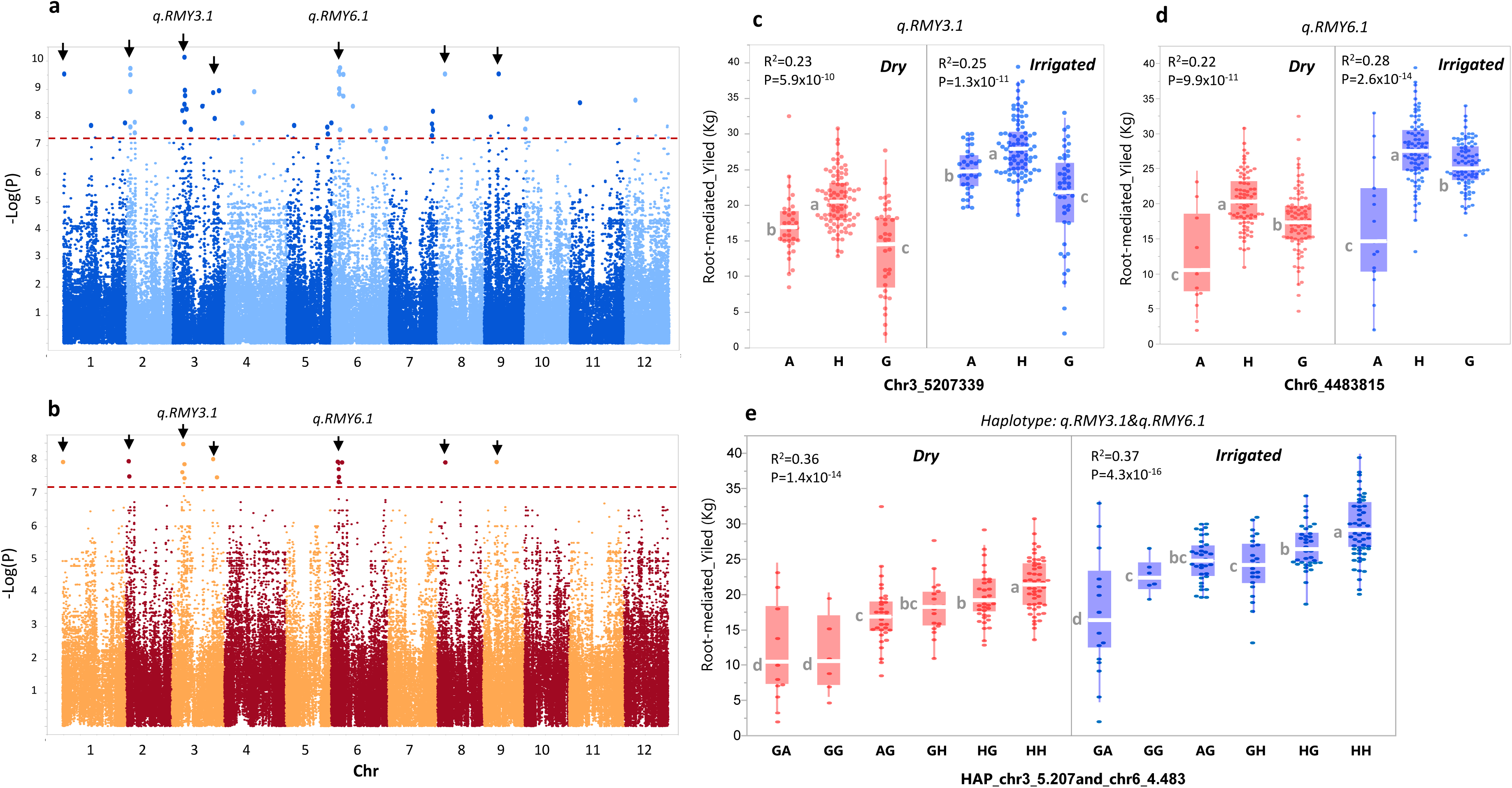
GWAS of root-mediated yield across 190 *HDA20* hybrids. a) Manhattan plot, Irrigated field. b) Manhattan plot, Dry field. Arrows indicate significant SNPs that are common to the Irrigated and Dry experiments. c) ANOVA for allelic effect of QTL on chromosome 3 (qRMY3.1). d) ANOVA for allelic effect of QTL on chromosome 6 (qRMY6.1). e) ANOVA for allelic effect of the combined haplotype of qRMY3.1 and qRMY6.1.

### Validation of selected hybrid rootstocks with multiple scions

Based on the large-scale analysis of rootstocks performance under two environments, we were able to select four high-yielding hybrid rootstocks for further testing. Scion x rootstock interactions are common in grafted plants and therefore, we grafted the selected rootstocks with four scions that represent different melon variety types: ‘Glory’ – reticulatus, long shelf-life Galia type; ‘Noy-Amid’ – inodorous, yellow canary type; ‘Hudson’ - reticulatus, ‘Ananas’ type and ‘HDA005’ - an experimental small-fruited (300 g) inter sub-specific hybrid. The four scion varieties were used as non-grafted controls in addition to two other control rootstocks: ‘Dulce’ - a *reticulatus* inbred line and one of the parents in the *HDA20* set, and ‘Tatsacabuto’, an inter-specific *Cucurbita* hybrid rootstock used commercially in melon and watermelon fields. **Figure 8a** is summarizing the results of the 28 scion x rootstock combinations from multiple field experiments representing different locations, planting densities and irrigation regimes. Yield performance of the different combinations is presented as percentage difference from the corresponding non-grafted scion variety; in a unified analysis of this experiment, the selected hybrid rootstocks significantly increased yield compared to the control varieties by 11% to 19% (**Figure 8a**, unified mean). While interactions between rootstock and scion and between genotype and environment existed across the different combinations, we find a significant overall yield advantage mediated by our selected experimental hybrid rootstocks over the commercial *Cucurbita* rootstock and the corresponding non-grafted scion varieties. We further tested two selected hybrid rootstocks the next year under two scions (‘Glory and ‘Noy-Amid’) and in two irrigation regimes and two planting densities (**Figure 8b**). The advantage of our experimental hybrids over the control rootstocks and self-grafted varieties was consistent in both scions and more prominent under standard planting density compared to wide spacing. These results that are based on yield analysis of more than 4,500 grafted plants over the different experiments conducted with the selected rootstocks in both years, provide an important proof-of-concept for the potential of hybrid rootstocks as a possible alternative channel for yield improvement in melon.

**Figure 8:**
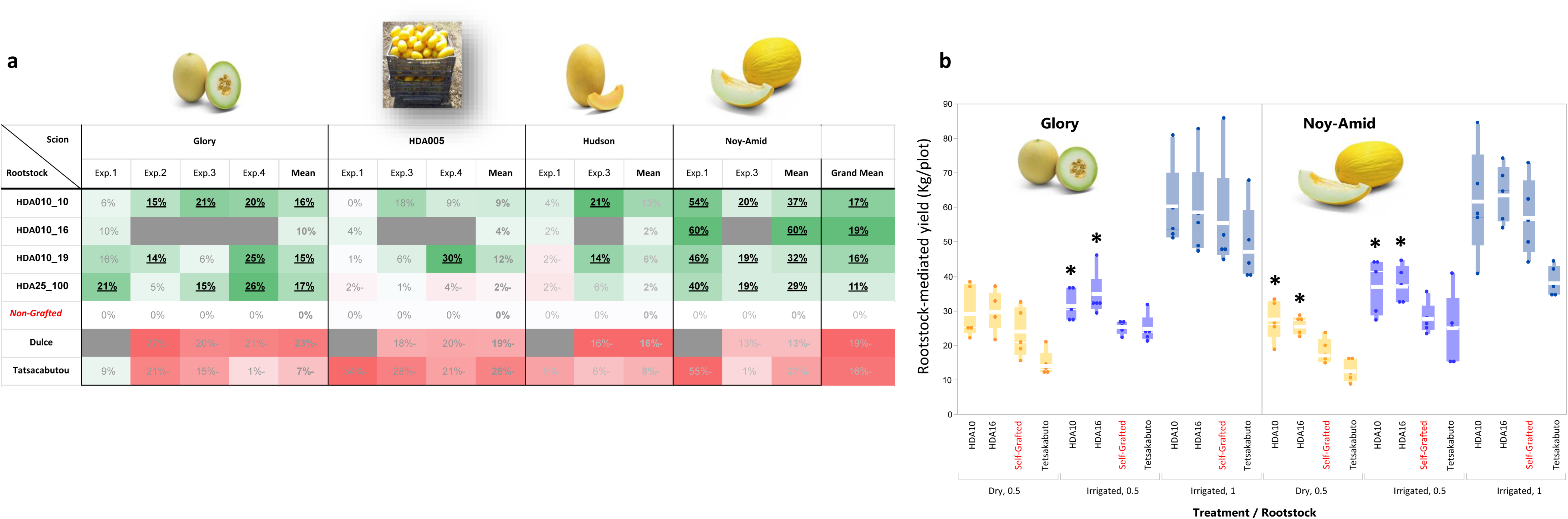
Yield advantage of selected rootstocks across scions and growing conditions (a) 2019 yield trials. Values in each cell are the average of 5 plots with 10 plants per plot and are presented as Δ% from the corresponding non-grafted variety. Significant values at P<0.05 are bolded and underlined. EXp.1: Maoz-Haim, Irrigated, 1.66 pl./m.; Exp.2: Newe-Ya’ar, Dry, 2 pl./m.; Exp.3: Newe-Ya’ar, Irrigated, 2 pl./m.; Exp.4: Newe-Ya’ar, Irrigated, 1 pl./m. (b) 2020 yield trials. * indicate significantly different (at P<0.05) from the self-grafted controls.

## Discussion

### Fruit yield heterosis in melon is prevalent and controlled independently above and underground

Charles Darwin noted already in 1876 that cross-pollinated F1 hybrids are more vigorous and productive than their parents (Darwin, 1876). Hybrid vigor, later termed heterosis to discriminate it from heterozygosity (Shull, 1948), is still intriguing geneticists and is commonly utilized for crop improvement (Duvick, 2001; Hochholdinger and Baldauf, 2018). While yield heterosis was extensively described in multiple plant species, so far it was investigated in a limited number of studies in melon, with variable conclusions regarding its magnitude and breeding impact (Katherine *et al.*, 2011; Pouyesh *et al.*, 2017; Napolitano *et al.*, 2020). In the current study, we initially show that as in other self and cross-pollinated crop plants, there is substantial yield heterosis also in melon. The average yield of the 45 diallel hybrids from our *HDA10* population was 73% higher compared to the average of their parents and almost 1/3 of these hybrids displayed significant BPH (**Figure 2a, b**). The yield heterosis was explained by combined effects on fruit number, average fruit weight and the tradeoff between them. An inherent drawback of studying yield heterosis across such diverse multi-parental melon population lay in the fact that the yield variation is potentially confounded by substantial variation in other morphological and developmental traits across the diversity. For example, variation in female sex expression type (monoecious or andromonoecious, (Gur *et al.*, 2017), 50-fold fruit weight variation (60-3500 g, **Figure 5b**) or substantial variation in earliness (85-120 days to maturity) were characterized across our population. These effects expand the overall phenotypic variation for multiplicative trait such as yield, and complicate the interpretation of genetic analyses. To dissect yield heterosis more effectively, we therefore took advantage of the fact that melon is amenable for grafting and allows physical separation and re-assembly of roots and shoot combinations. We focused our yield analysis on root-mediated effects by performing a common-scion rootstock experiments. While, as expected, the overall coefficient of variation (CV) of yield in the common-scion grafted experiment was less than a third of yield CV in the parallel non-grafted experiment (0.29 and 1.02, respectively), the broad sense heritability was very similar (H^2^=~0.40), confirming the effectiveness of this approach and the significant heritable contribution of roots to yield variation. We detected prominent yield heterosis both above (non-grafted) and underground (root-mediated), but the correlation between these setups was low (**Figure 2f**), which makes sense considering the substantial morphological and physiological aboveground variation that is only partly dependent on roots function, and the probable cross talk between root and shoot. The significant root-mediated effects that we describe here for yield variation and heterosis emphasize the essential, underestimated, contribution of roots to whole plant phenotype. It is important to note, however, that root-mediated effects were not common to all traits. For example, rind netting or internal and external color of ‘Glory’ fruits did not display notable visual differences across the 210 different rootstocks (data not shown). Another quantitative example for that is fruit TSS, for which we find substantial heritable variation across the 210 *HDA20* genotypes in non-grafted plants (3%-16% Brix) but minor, non-significant, root-mediated effects in the common scion experiments (**Sup. Figure 5**). This indicates that fruit TSS is determined largely by above-ground (canopy) properties, including genetically controlled fruit metabolism (Burger and Schaffer, 2007).

### Root-mediated yield variation is positively correlated with variation in both Fruit Number (FN) and Average Fruit Weight (AFW)

Analysis of yield components across more than 7,300 common-scion grafted rootstocks in the multi-allelic *HDA20* population revealed 3-fold range for FN and 1.7 fold for AFW (**Sup. Figure 3a-d**) with significant positive correlations of both traits with yield, and accordingly also positive correlation between these two components (**Sup. Figure 3e, f**). This pattern is in complete contrast to the significant negative tradeoff observed between AFW and FN across our non-grafted melon diversity, where increase in AFW is strongly associated with decrease in FN (R^2^=0.75, **Sup. Figure 4**). Tradeoff between yield components is a common pattern in plants (Nesbitt *et al.*, 2001; Golan *et al.*, 2019; Gadri *et al.*, 2020) and may reflect evolution of developmental plasticity that promote reproductive fitness stability. More generally, trade-off between size and number is common across biological systems and can be explained simply as a result of limited resources (Garland, 2014). The absence of negative tradeoff between AFW and FN in our rootstock experiments, expressed as parallel increase in both FN and AFW in high-yielding rootstocks, suggest that the rootstocks variation is associated with modifications in resources availability or in alterations of sink-source relations in a way that is not interfering with the developmental program of the scion genotype.

### Mode of inheritance of reproductive vs. morphological or metabolic traits in melon

We show here that ‘Underground’ yield heterosis is a prominent attribute in melon (**Figures 2d, e Figure 3**) and that most of the root-mediated yield variation across 190 diverse *HDA20* hybrids can be explained by non-additive genetic components (**Figure 4**). Comparisons to the mode of inheritance of AFW, ASW and TSS, measured on non-grafted plants across the same *HDA20* population (**Figure 5**), indicates that heterosis in melon is more prevalent in reproductive traits compared to non-reproductive (morphological or metabolic) traits. This observation confirms the similar phenomena previously described in maize (Flint-Garcia *et al.*, 2009), tomato (Semel *et al.*, 2006) and mice (Rocha *et al.*, 2004). This fundamental difference in mode-of-inheritance between trait categories, that is consistent across diverse taxonomic groups, indicates a possible evolutionary role of this pattern. Our results expand the perspective on this, as we show here that even the exclusive effect of roots variation on whole-plant performance, maintain the prominent heterotic mode-of inheritance of total fruit yield and canopy biomass across natural melon diversity.

### Prediction of root-mediated yield heterosis

Heterosis, the positive deviation of hybrid from its parental mean is at the same time desired and challenging genetic property for plant breeders. Predicting and maximizing heterotic response in F1 hybrids is a challenge, as parental performance *per se* are not necessarily informative. The development of prediction tools or breeding strategies to maximize the chances for producing successful crosses is therefore a key objective in hybrid breeding (Bernardo, 1994; Zhao *et al.*, 2015). We show here that root-mediated yield of melon hybrids is superior, but independent of their parental *per se* performance (**Figure 4, Figure 5a**), and therefore implementation of high-throughput indirect selection or prediction methods is important for efficient rootstock breeding. Root-mediated early-stage vegetative canopy biomass was not predictive as a potential indirect selection trait. Parental genetic distance was also poorly correlated with root-mediated hybrids yield. However, our GWA results (**Figure 7**) indicate that QTL or genomic selection strategies can be effective for accelerating rootstock breeding. Haplotype of two QTLs that were consistent across Irrigated and Dry experiments, explained 36% of the root-mediated yield variation and the favorable haplotype (heterozygote at both loci) was associated with average yield increase of 15% compared to the *HDA20* population mean.

### Breeding implications

World population growth and global climate change are forming major challenges to our civilization (Godfray *et al.*, 2010; Wheeler and von Braun, 2013). Agriculture, among other disciplines, plays a key role in dealing with these challenges (Garnett *et al.*, 2013) and one of the important channels of action for improving yields of crop plants in a sustainable manner is through genetic research and breeding. Heterosis is a well-established genetic mechanism for yield enhancement in crop plants. While parental genetic distance *per se* is not necessarily a robust predictor for level of heterosis in F1 hybrids—as shown here and by others (Huang *et al.*, 2015; Yang *et al.*, 2017b; Kaushik *et al.*, 2018) —it is a consensus that stronger heterotic effects are expected in hybrids by crossing diverse rather than closely related parents. Commercial melon breeding is commonly performed within market-segment defined narrow germplasm pools, which on one hand ensures strict maintenance of fruit-related varietal characteristics, but on the other hand inhibits the ability to perform wide crosses and explore the full potential of heterosis for productivity traits. By focusing our yield enhancement research effort on rootstocks, we essentially bypass this barrier as the above and underground genetic actions are performed independently. We show here that melon hybrid rootstocks significantly outperform inbreds and that selected melon hybrids, used as rootstocks grafted with commercial melon variety, increase yield across scions and environments without any visible negative effect on fruit quality. The ability to implement focused and autonomous breeding for rootstocks to efficiently introduce beneficial genetic properties to roots in species amenable for grafting, is a powerful, currently underutilized approach to improve crop performance under optimal as well as stress conditions. Mapping root-mediated heterotic yield QTLs in a multi-allelic population is a first step towards focused QTL analysis in bi-parental populations and development of marker-assisted selection protocols. Using hybrid-breeding methodologies, rootstock breeding can be an effective alternative channel for development of stress-tolerant and high-yielding varieties in crop species that are suitable for grafting, such as *Cucurbitacea* and *Solanaceae*.

### Inverted scheme in root genetics

Root biology is receiving increased attention in recent years as a potential channel to improve plant productivity under optimal and stress conditions. However, most of the genetic research in model and crop plants is taking an inherent approach with initial focus on analysis of root development and variation in root system architecture (Bray and Topp, 2018; Zhao *et al.*, 2018; Jia *et al.*, 2019; Wachsman *et al.*, 2020), rather than direct analysis of roots functional variation. Here, we propose an inverted scheme; using grafting, we directly characterize variation in root function and effect on whole-plant performance in the field to study the genetics of root-mediated yield variation. The combination of a crop plant amenable for grafting, with rich genetic and genomic resources, such as melon, is a powerful platform for applied root genomics and for exploring the interactions between root and shoot. We, therefore, believe that such ‘forward genetics’ approach is a first step towards discovery of candidate genes involved in root function, that show proven effect on yield. The current research expands the view on genetic properties of heterosis in plants by highlighting the contribution of roots to yield heterosis.

## Supplementary data

**Supplementary Table 1:** List of 25 Founder lines that compose the melon core subset.

**Supplementary Figure 1:** Structure of the Half-Diallel (HDA) sets.

**Supplementary Figure 2:** *HDA20* rootstock yield trials in summer 2018 (Irrigated and Dry). a) Grafted plants in the nursery just before transplanting. Plastic clips are the graft union positions. b) Our field at Newe Ya’ar during yield harvest. Melon piles are the yield of plots of five plants. c) Yield heatmap projected on the 1,462 field plots (7,310 plants) of the Dry and Irrigated experiments. d) Correlation between Dry and Irrigated trials. Each dot represents an entry mean in the Dry and Irrigated fields. e and f) Correlations between root-mediated yield in 2017 and 2018 (irrigated and Dry) across 55 *HDA10* genotypes. The common scion, ‘Glory’, grafted on itself (Gr) and non-grafted (NG) are highlighted.

**Supplementary Figure 3:** Correlations between root-mediated yield and its components – Number of Fruits per plant (FN) and Average Fruit Weight (AFW), in the *HDA20* population in the Irrigated and Dry fields.

**Supplementary Figure 4:** Correlation between Average Fruit Weight (AFW) and Fruit Number (FN) across 45 *HDA10* F1 hybrids and their 10 parents. a) Normal scale. b) Log transformed values

**Supplementary Figure 5:** Correlation for TSS between the rootstock-mediated ‘Glory’ and non-grafted experiments across the *HDA20* population. Each point represent the entry mean TSS of 15 fruits in the grafted (rootstock-mediated, x-axis) and non-grafted (y-axis) experiments.

## Acknowledgments

We wish to thank Uzi Saar, Fabian Baumkoler and the Newe-Yaar farm team for technical assistance in setting the field trials and for plant maintenance. We thank Jeff Glaubitz and the team at the Genomic Diversity Facility at Cornell University for sequencing services. Funding for this research was provided by the United States-Israel Binational Agricultural Research and Development Fund (BARD) grant no. IS-4911-16 and by the Israel Science Foundation (ISF) grant no. 860/19.

## Author contributions

AG conceived the research plan. AG and AD designed the experiments. AD, JB, and AG developed plant genetic materials. AD, IH, EO, GT, AM, TI and AG performed the experiments and collected the data. AG and AD analyzed the results. AAS, YT and ESB provided genomic support. AG wrote the manuscript. All authors discussed the results and approved the manuscript.

