## Supplementary figures and images for "Underground Heterosis for Melons Yield"

### Supplemental Figure 1

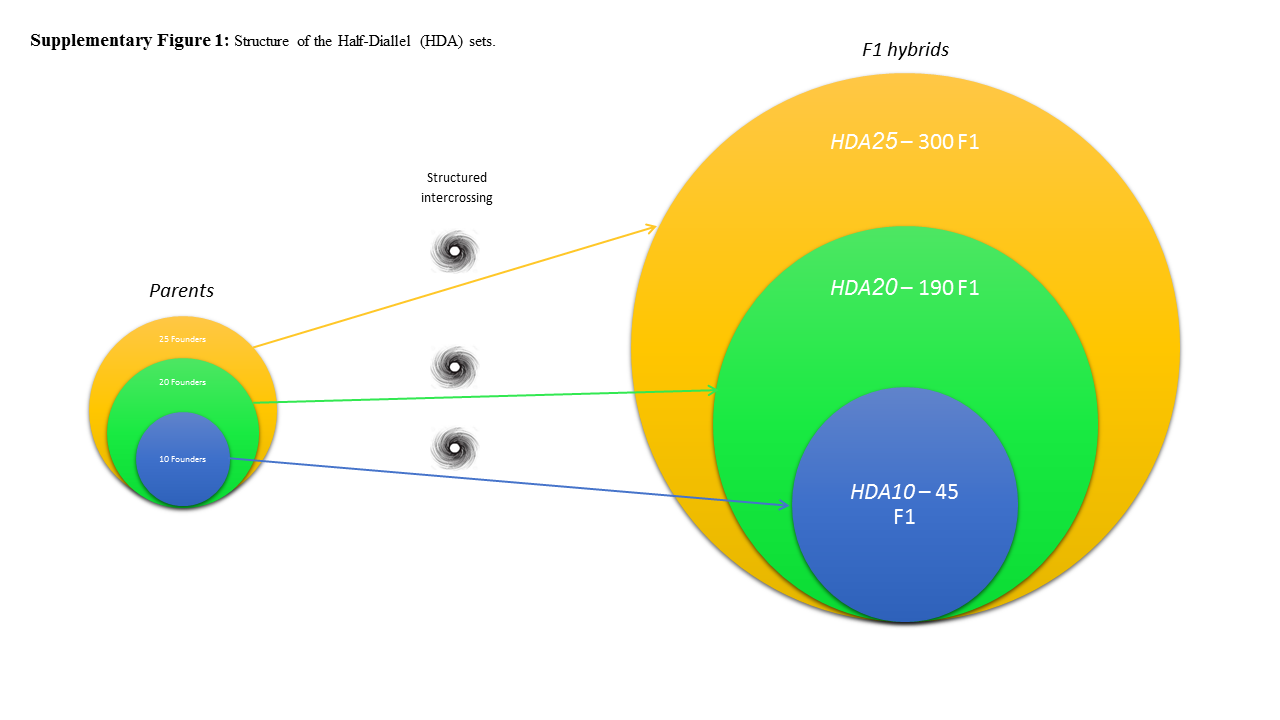

### Supplemental Figure 2

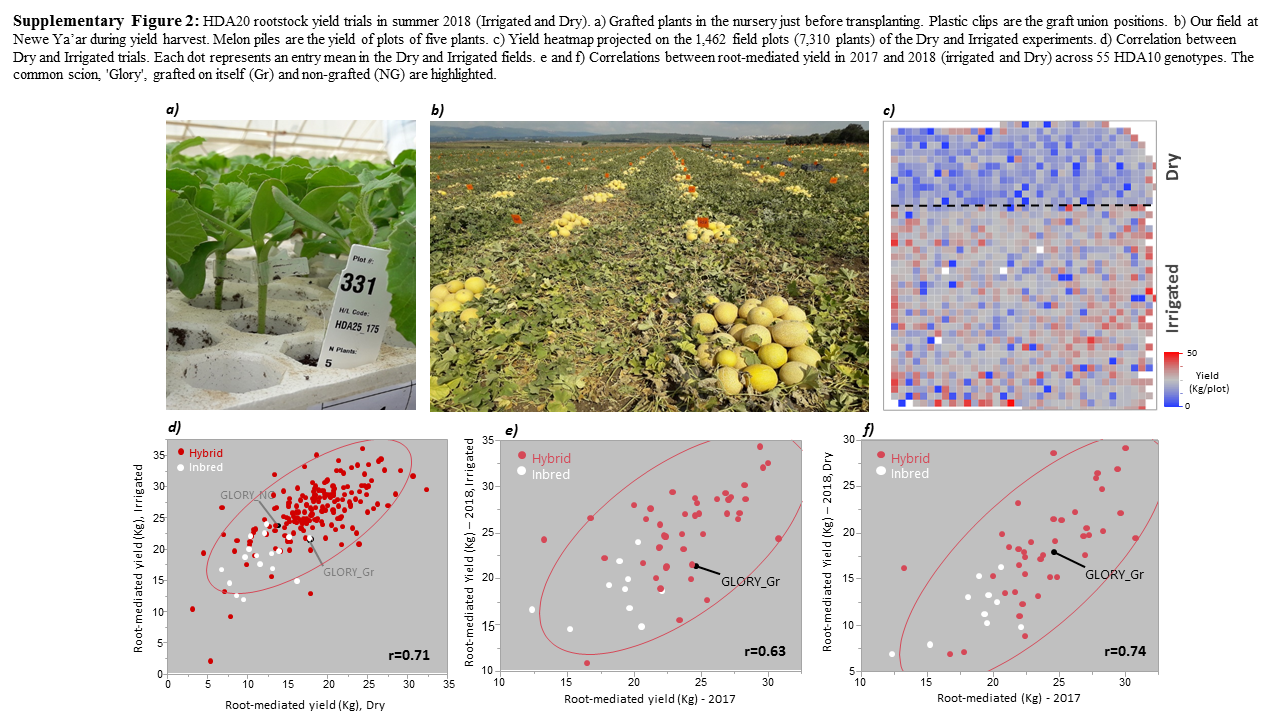

### Supplemental Figure 3

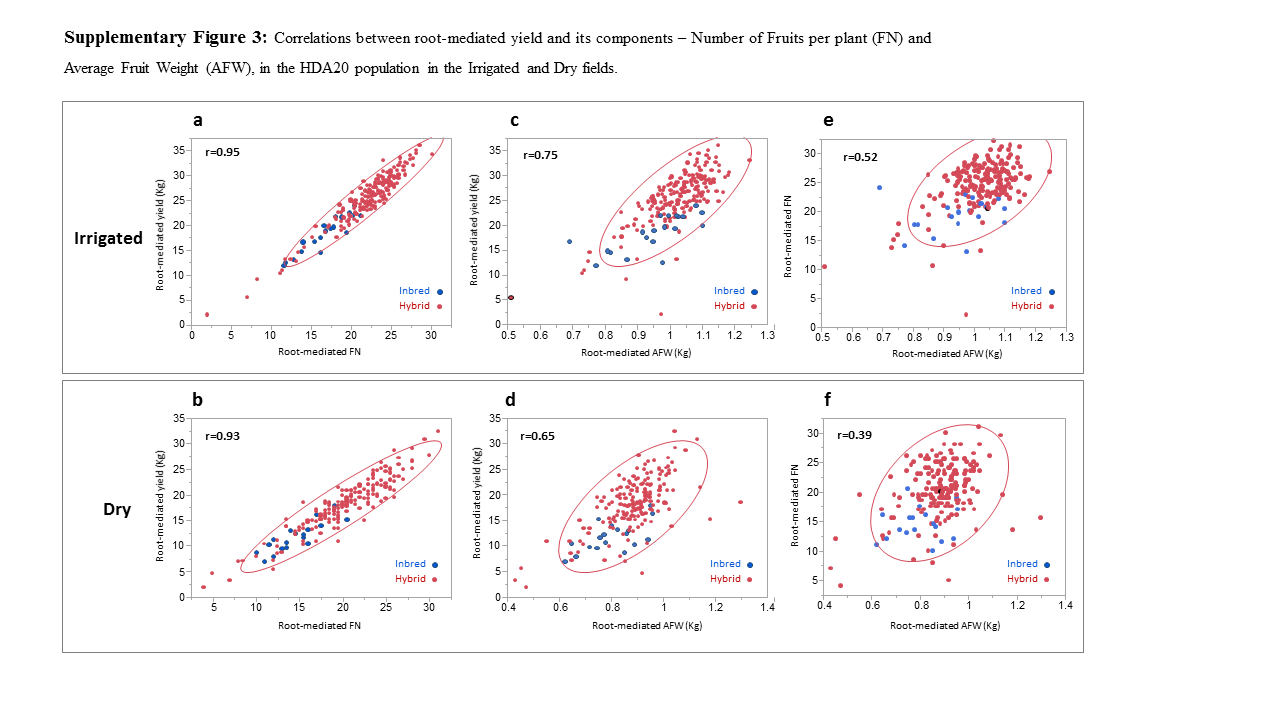

### Supplemental Figure 4

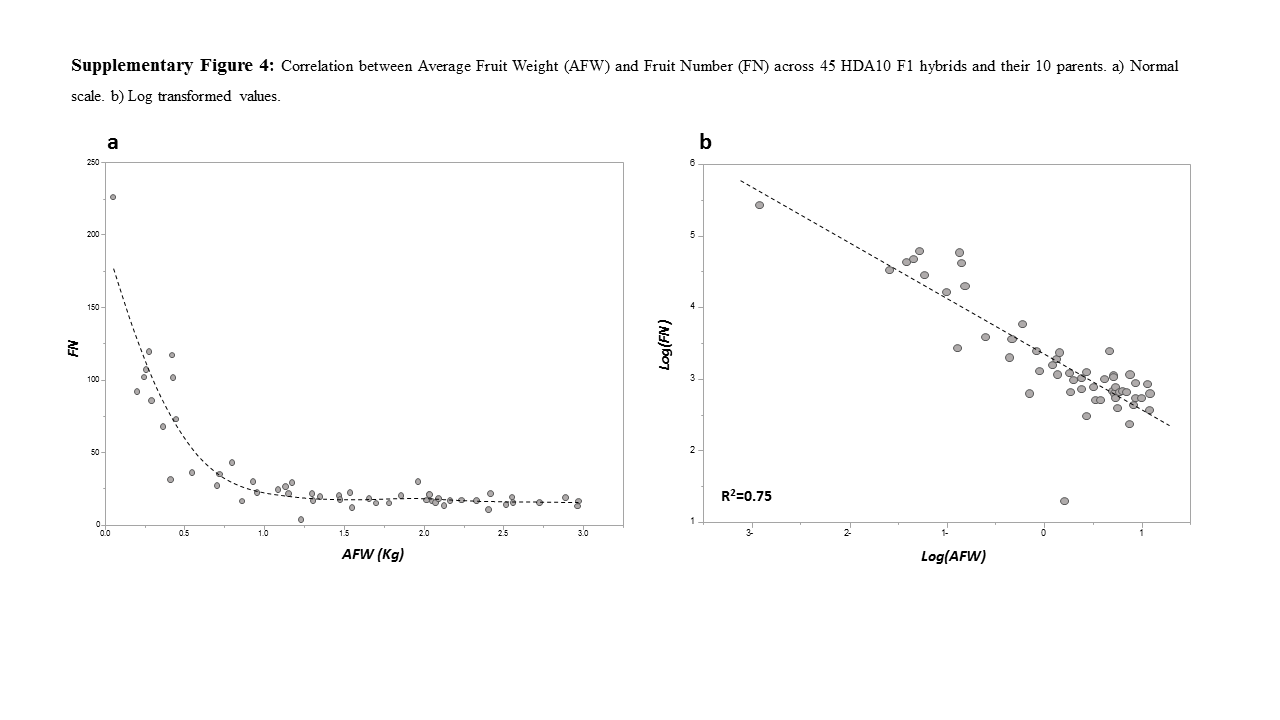

### Supplemental Figure 5

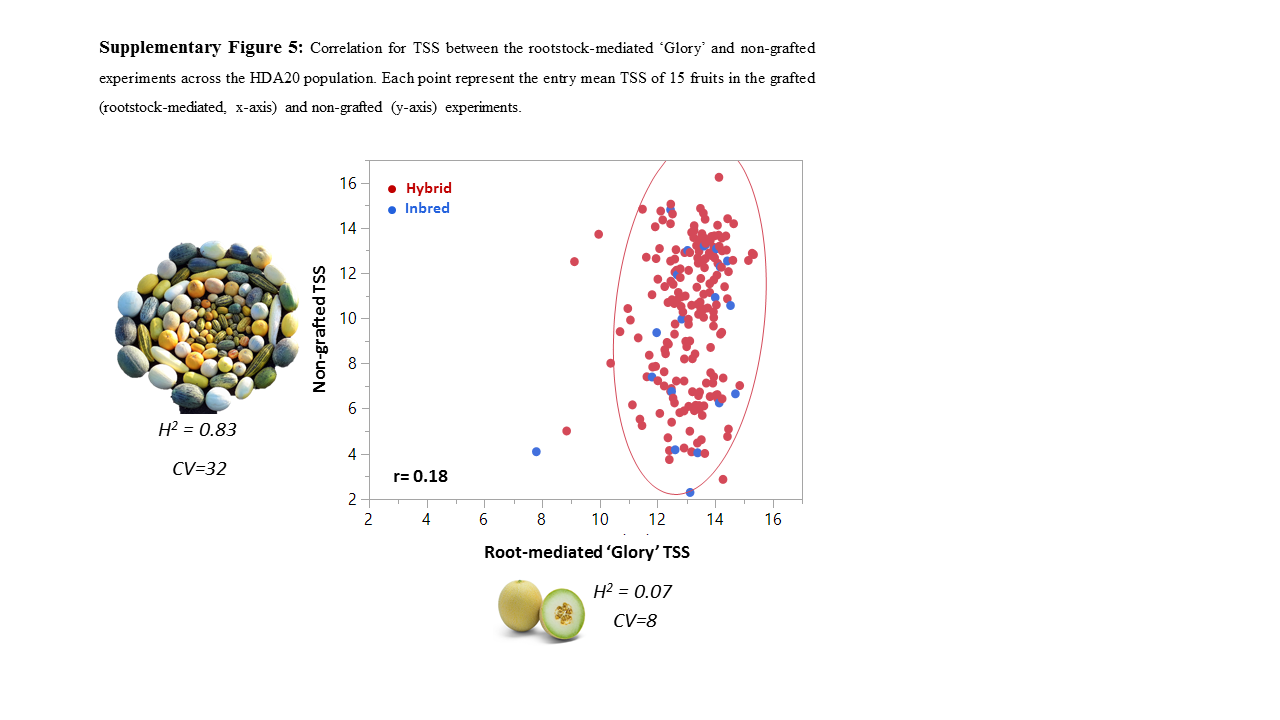

### Supplemental Table 1

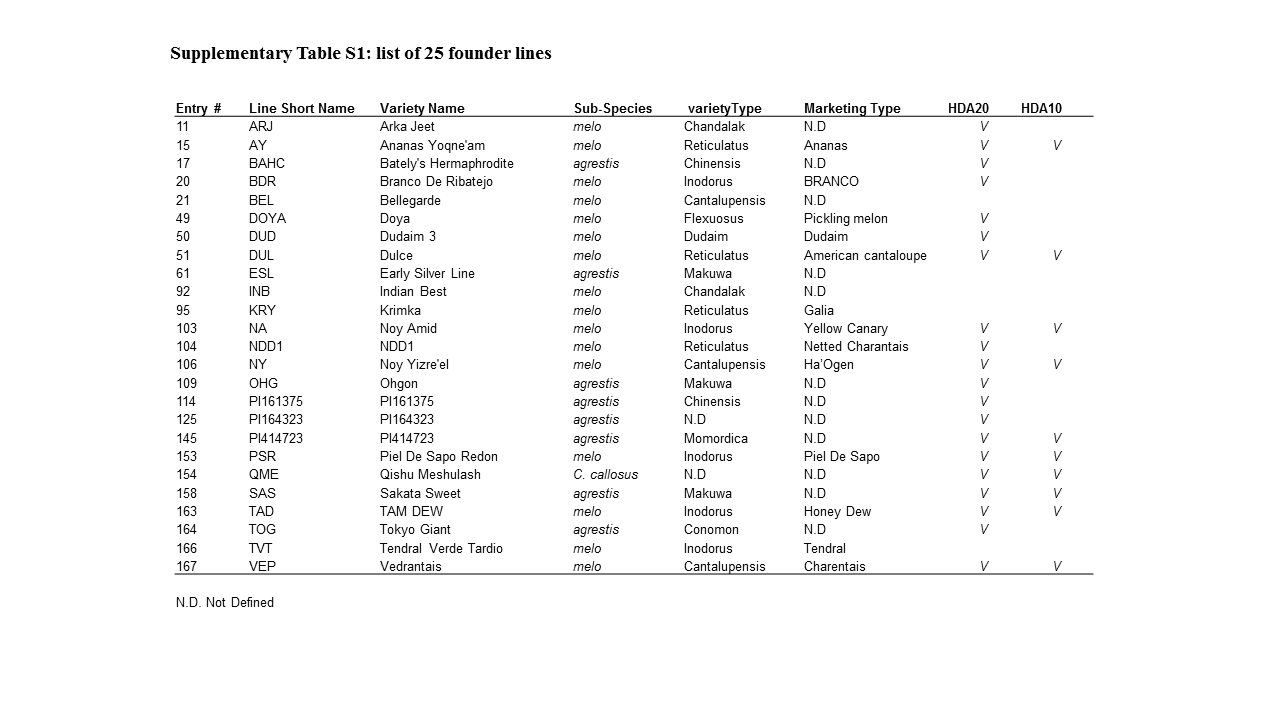
